# Type I PRMTs Play a Role in Mammalian Embryonic Lineage Specification

**DOI:** 10.64898/2026.08.18.745309

**Authors:** Jinlong Qiu, Yutong Chen, Pedro Beltran-Alvarez, Roger Sturmey

## Abstract

Mammalian preimplantation development requires precisely coordinated lineage decisions to establish the trophectoderm (TE), inner cell mass (ICM), epiblast (EPI), and primitive endoderm (PrE). Glucose metabolism and epigenetic regulation are increasingly recognised as key determinants of lineage specification during preimplantation development. However, how glucose-dependent metabolic cues interface with epigenetic mechanisms to regulate embryonic cell fate remains poorly understood. Here, we investigated the role of glucose in regulating protein methylation by protein arginine methyltransferases (PRMT) in bovine preimplantation development. PRMT1 and its associated histone mark H4R3me2a were detected throughout bovine oocyte maturation and embryo development. Pharmacological inhibition of Type I PRMTs using two structurally distinct inhibitors, GSK3368715 and MS023, markedly reduced global protein asymmetric dimethylarginine (ADMA) and H4R3me2a levels. PRMT inhibition impaired blastocyst cell proliferation, reduced total cell number, and disrupted both first and second lineage decisions, as demonstrated by decreased CDX2- and SOX2-positive TE and ICM cells and reduced NANOG- and GATA6-positive EPI and PrE cell allocation. Mechanistically, Type I PRMT inhibition downregulated key components of the Hippo-associated TE programme, including YAP, TEAD4, and TFAP2C. Consistent effects were observed in mouse embryos, where MS023 treatment reduced ADMA, CDX2, YAP, and TFAP2C expression and impaired TE and ICM allocation. Collectively, our findings identify Type I PRMT-mediated ADMA as an essential epigenetic regulator of early mammalian lineage specification and support a conserved ADMA–Hippo regulatory axis linking arginine methylation to embryonic cell fate decisions.

**In brief:** Type I protein arginine methyltransferase (PRMT)-mediated asymmetric dimethylarginine (ADMA) is required for proper lineage specification during mammalian preimplantation development. ADMA depletion disrupts Hippo signalling, cell proliferation, and trophectoderm and inner cell mass allocation in bovine and mouse embryos.

## Introduction

Mammalian preimplantation embryonic development is a highly orchestrated process that culminates in two sequential cell fate decisions essential for successful implantation and fetal development [1, 2]. During the morula-to-blastocyst transition, the first cell fate decision establishes the trophectoderm (TE), which gives rise to the placenta, and the pluripotent inner cell mass (ICM), which forms the embryo proper [1, 2]. Subsequently, the ICM undergoes a second cell fate decision to generate the pluripotent epiblast (EPI), which gives rise to the fetus, and the primitive endoderm (PrE; hypoblast in bovines), which contributes to the yolk sac [1, 2]. These lineage segregation events are orchestrated by mutually antagonistic transcriptional networks, including CDX2 for TE specification, SOX2 for ICM identity, and NANOG versus GATA6 for EPI and PrE specification [1, 2].

Recent studies have highlighted the emerging roles of metabolic cues and epigenetic regulation in shaping lineage specification during preimplantation development. In mice, glucose promotes TE differentiation by modulating Hippo pathway activity, thereby facilitating nuclear YAP1 accumulation and formation of the YAP1–TEAD4– TFAP2C transcriptional complex, which activates CDX2 expression [3]. We recently demonstrated that this glucose–Hippo–YAP1 regulatory mechanism is conserved in bovine embryos [4]. A previous metabolomic study of bovine preimplantation embryos revealed that glucose availability markedly alters the intracellular abundance of symmetric and asymmetric dimethylarginine (SDMA and ADMA) [5]. Consistent with this observation, our preliminary immunofluorescence (IF) analysis showed that glucose deprivation substantially reduces global ADMA levels in cattle embryos [4]. However, the roles of ADMA and SDMA in bovine preimplantation embryos remain unknown, limiting our understanding of whether glucose regulates lineage specification through ADMA- and SDMA-associated metabolic or epigenetic mechanisms.

ADMA is generated by type I protein arginine methyltransferases (PRMTs), which in mammals comprise PRMT1, PRMT2, PRMT3, PRMT4, PRMT6, and PRMT8 [6]. During the first cell fate decision, PRMT4 catalyses histone arginine asymmetrical dimethylation to promote the expression of pluripotency-associated genes, including SOX2, SOX21, and NANOG, while repressing CDX2, thereby favouring inner cell mass specification [7-9].

During the second cell fate decision, PRMT1 methylates the pluripotency factor KLF4, facilitating recruitment of the mSin3a/HDAC co-repressor complex to repress GATA6 and SOX17, thereby restricting PrE differentiation and promoting epiblast identity [10]. Collectively, these studies indicate that type I PRMTs play critical roles in regulating lineage specification during mammalian preimplantation development; however, their functions and underlying mechanisms in bovine embryos remain largely unknown.

We therefore investigated the role of type I PRMTs mediated ADMA during bovine preimplantation development. Our study demonstrates that reduced ADMA levels markedly impair Hippo signaling activity in both mouse and bovine embryos, leading to decreased CDX2 expression. In bovine embryos, inhibition of type I PRMTs also disrupts the second cell fate decision. Collectively, our findings indicate that type I PRMT-mediated ADMA regulation is essential for proper lineage specification during bovine preimplantation development, linking glucose metabolism to Hippo signaling and epigenetic control of cell fate.

## Materials and Methods

### Reagents and Ethical Approval

Unless otherwise specified, fundamental chemical reagents utilized throughout this study were procured from Sigma-Aldrich. All animal handling and experimental procedures were conducted in strict accordance with the ethical guidelines and were officially sanctioned by the University of Manchester (Project License PD7C22AA9).

### Animal Models and In Vitro Embryo Culture

Mouse Embryo Collection: CD-1® IGS mouse were purchased from Charles River UK (Kent, UK). Superovulation was induced in 8-to-10-week-old female mice via I.P. injection of 5 IU pregnant mare’s serum gonadotrophin (PMSG, Intervet®, UK), followed by 5 IU human chorionic gonadotrophin (hCG, Chorulon®, UK) 48 hours later, prior to mating with fertile males. Zygotes were harvested 18–20 hours post-hCG injection, mechanically denuded in M2 medium with hyaluronidase (#MR-051), and maintained in KSOM medium (#MR-106) at 37°C in a humidified incubator with 5% O2 and 5% CO2 supply.

### Bovine In Vitro Production (IVP)

Bovine ovaries were sourced from a local slaughterhouse in Yorkshire, UK. Cumulus-oocyte complexes (COCs) presenting multiple cumulus cell layers were matured in vitro for 22–24 hours at 38.5°C in BO-IVM media (IVF Bioscience). Following maturation, COCs were co-incubated with BO-SemenPrep (IVF Bioscience)-purified spermatozoa (1–5 × 10^6^) in BO-IVF medium (IVF Bioscience) for 22 hours. Presumptive zygotes were subsequently denuded and cultured in BO-IVC medium (IVF Bioscience) under mineral oil (IVF Bioscience) at 38.5°C in a 5% CO2 and 5% O2 atmosphere until further evaluation.

### Inhibitor Treatment

To evaluate the functional consequences of targeted PRMT inhibition, preimplantation embryos (both mouse and bovine) were treated with the GSK3368715 or MS023 commencing at the zygote stage. The stock solution of the inhibitor was prepared by dissolving the compound in dimethyl sulfoxide (DMSO). The inhibitor was subsequently supplemented into the respective culture media—BO-IVC medium for bovine embryos and KSOM medium for mouse embryos. For both species, the final working concentration of DMSO in the culture media was strictly maintained at 2.8 mM in both the inhibitor-treated and vehicle control groups to rule out solvent-induced effects.

### Immunofluorescence and Confocal Microscopy

Cultured embryos were fixed in 4% paraformaldehyde for 10–30 minutes, permeabilized with 0.5% Triton X-100 in PBS, and subjected to a blocking step utilizing 10% FBS combined with 0.1% Triton X-100. The specimens were then incubated with primary antibodies overnight at 4°C, followed by a 2-hour incubation with corresponding secondary antibodies at room temperature. Nuclei were counterstained with Hoechst3342 (Hoechst) for bovine and DAPI for mouse. Fluorescence imaging was conducted utilizing either a standard epifluorescence microscope or a Zeiss LSM710 (cattle)/ Leica SP8x (mouse) confocal laser scanning microscope, with Z-stack projections generated via maximum intensity algorithms. Signal quantification was performed using ImageJ software by calculating the specific nuclear fluorescence intensity after subtracting the corresponding cytoplasmic background signals. Specific antibody details can be referenced in Supplementary Table S1

### Statistical Analysis

Quantitative data were processed and visualized using GraphPad Prism 10.0. Differences between two experimental groups were assessed via two-tailed unpaired Student’s t-tests. For multigroup comparisons, a one-way analysis of variance (ANOVA) followed by Tukey’s post-hoc test was applied. All experiments were performed with a minimum of three independent biological replicates, and statistical significance was defined as P < 0.05.

## Results

Given that PRMT1 is established as the predominant enzyme responsible for about 85% of cellular ADMA synthesis [6], we sought to further characterize its mRNA and protein-level dynamics. We first analyzed public RNA-seq datasets [11] to generate the mRNA profiles of PRMT1 throughout cattle oocyte maturation and preimplantation embryonic development. Our analysis revealed dynamic expression patterns of type I PRMTs from the germinal vesicle (GV) stage to the blastocyst (BL) stage; however, no significant differences were detected between adjacent developmental stages (Fig. 1A).

**Fig. 1.**
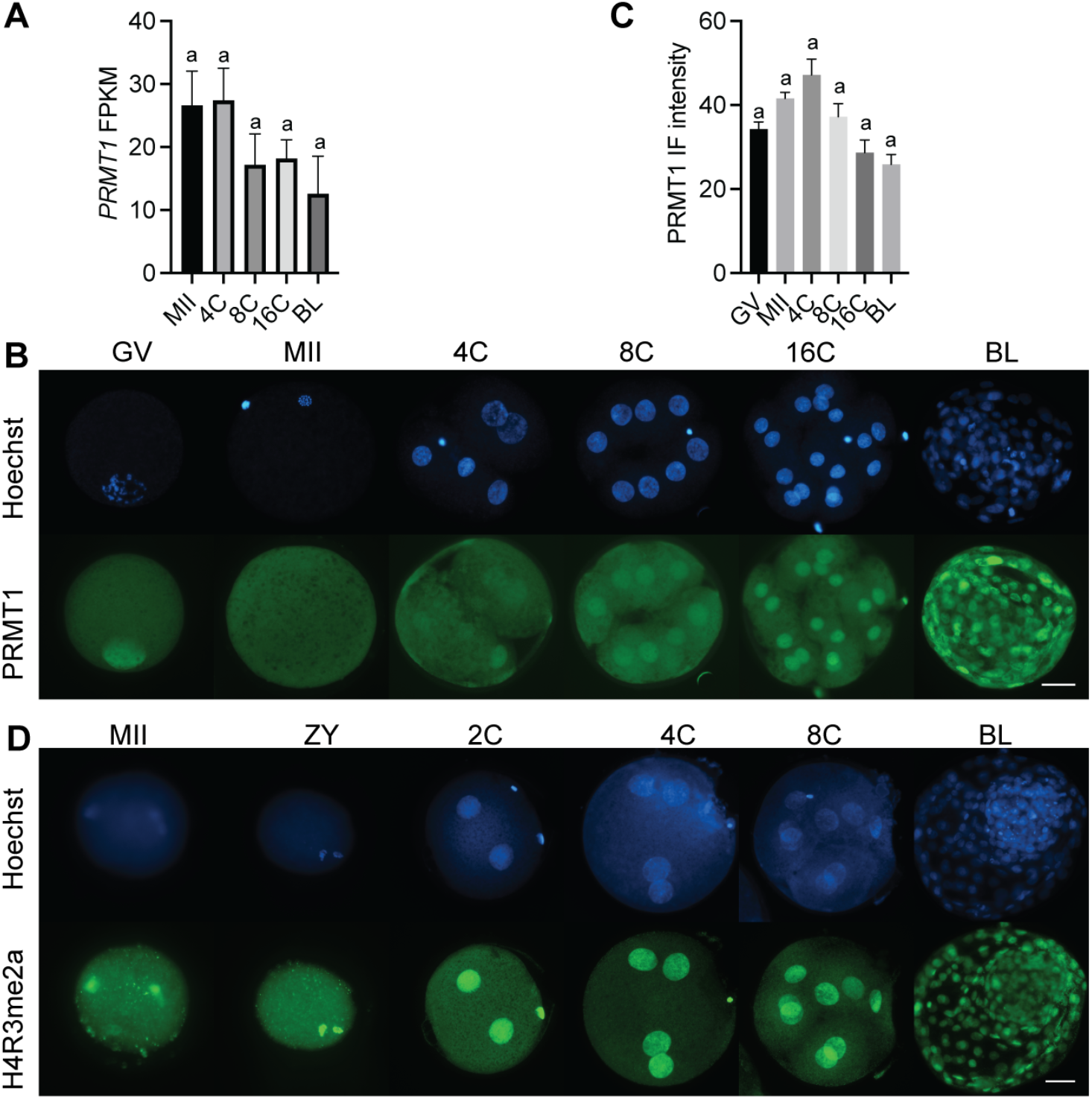
Dynamic mRNA and protein profiles of PRMT1 and H4R3me2a during preimplantation development in cattle. (A) *PRMT1* mRNA profiles in cattle generated using public RNA-seq datasets. (B, C) IF detection of PRMT1 protein during oocyte maturation and embryonic development in cattle. (D) IF detection of H4R3me2a during oocyte maturation and embryonic development in cattle. GV: germinal vesicle, MII: metaphase II, Zy: zygote, 2C: 2-cell, 4C: 4-cell, 8C: 8-cell,16C: 16-cell, MO: morula, BL: blastocyst. n = 3 replicates. 15-20 oocytes/embryos were analysed at each stage. Data are presented as mean ± SEM. Values with different superscripts across different concentrations indicate significant differences (P < 0.05). Scale bar=100 µm.

Subsequently, using immunofluorescence (IF), we systematically detected the spatio-temporal protein distribution of PRMT1, alongside its specifically catalyzed histone modification mark H4R3me2a, throughout bovine oocyte maturation and preimplantation embryonic development (Fig. 1C, D). IF analysis revealed that PRMT1 is consistently expressed throughout preimplantation development across both the cytoplasm and the nucleus from the GV to the BL stages (Fig. 1C). Consistent with its mRNA expression profile, no significant differences were observed between adjacent developmental stages (Fig. 1D). H4R3me2a exhibited a strict nuclear localization from the MII stage through the blastocyst stage (Fig. 1E).

Collectively, these dynamic mRNA and protein profiles indicate that Type I PRMTs and their associated arginine methylation marks are actively established on the chromatin, strongly suggesting that they play a role in cattle preimplantation embryonic development.

### Inhibition of Type I PRMTs disrupts global arginine methylation levels

To investigate the role of type I PRMTs and ADMA in preimplantation development, we employed two highly selective, competitive type I PRMT inhibitors, GSK3368715 (GSK715) and MS023. The use of two structurally distinct inhibitors targeting the substrate-binding pocket minimizes the likelihood of off-target effects and strengthens confidence in any observed phenotypes [12, 13].

We first performed a dose-response analysis to identify inhibitor concentrations that effectively suppressed ADMA while avoiding non-specific toxicity or developmental arrest (Fig. 2A). Embryonic developmental competence was significantly compromised when embryos were exposed to GSK715 or MS023 concentrations higher than 20 μM or 5 μM, respectively (Fig. 2A-C). For this reason, further experiments were conducted using 10 and 1 μM GSK715 or MS023, respectively. IF analysis confirmed that 10 μM GSK715 and 1 μM MS023 effectively inhibited type I PRMT-mediated arginine methylation, as evidenced by markedly reduced H4R3me2a deposition (Fig. 2D, E) and global ADMA levels (Fig. 2F, G). These concentrations were therefore used in all subsequent experiments.

**Fig. 2.**
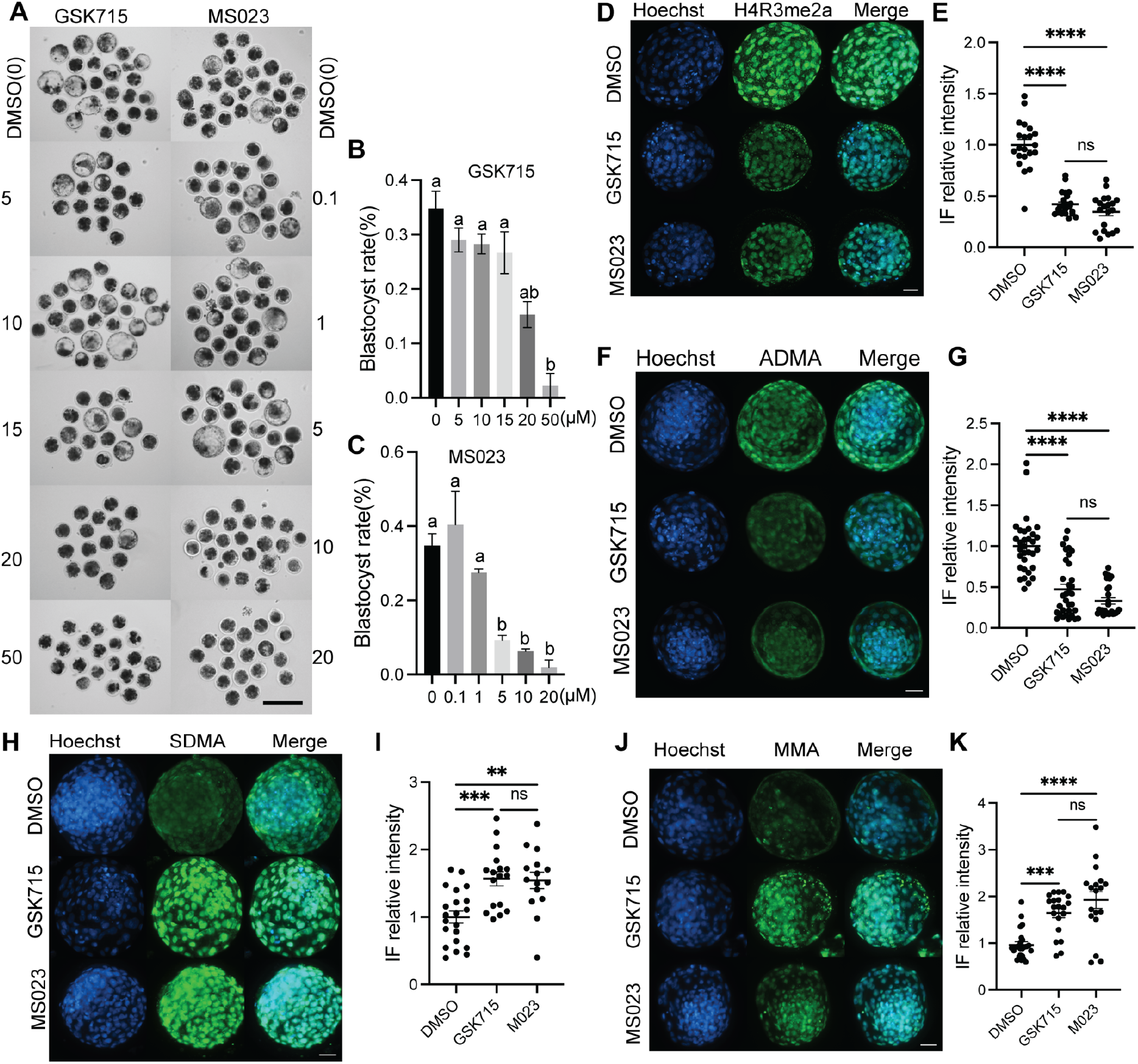
Inhibition of Type I PRMTs disrupts global arginine methylation levels. (A) Representative bright-field images of cattle blastocysts treated with various concentrations of GSK715 or MS023. Scale bar = 50 μm. (B, C) Blastocyst rates following treatment with di^erent concentrations of GSK715 (B) and MS023 (C). n = 3 replicates, with 15-25 embryos per group per replicate. Data are presented as mean ± SEM. Values with different superscripts across different concentrations indicate significant differences (P < 0.05). (D, F, H, J) Immunofluorescence analysis of H4R3me2a (D), ADMA (F), SDMA (H), and MMA (J) in blastocysts treated with DMSO, GSK715, or MS023. Scale bar = 100 μm. (E, G, I, K) Quantification of relative immunofluorescence (IF) intensity for H4R3me2a (E), ADMA (G), SDMA (I), and MMA (K) in DMSO, GSK715 and MS023 groups. Each dot represents an individual blastocyst. n = 3 replicates, with 5-12 blastocysts per group per replicate. Data are presented as mean ± SEM. Asterisks in indicate significant differences (**P < 0.01, ***P < 0.001, ****P < 0.0001; ns, not significant).

Given that type I, II and III PRMTs methylate overlapping arginine residues on protein substrates, inhibition of type I PRMTs is expected to increase substrate availability for type II and III enzymes [14]. Consistent with this mechanism, embryos treated with either GSK715 or MS023 exhibited a marked compensatory increase in symmetric dimethylarginine (SDMA) (Fig. 2H, I) and monomethylarginine (MMA) (Fig. 2J, K).

Together, these findings demonstrate that GSK715 and MS023 efficiently and specifically suppress type I PRMT-mediated arginine methylation in embryos and substantially remodel the global arginine methylation landscape during bovine preimplantation development.

### Inhibition of Type I PRMTs impairs lineage specification and cell allocation in cattle blastocysts

Having established conditions that effectively suppress Type I PRMT-mediated arginine methylation, we next examined whether ADMA depletion disrupts the two sequential cell fate decisions during cattle preimplantation development.

To assess the first cell fate decision, we analysed the expression of the TE marker CDX2 and the ICM marker SOX2. IF analysis showed that treatment with either GSK715 or MS023 significantly reduced the relative fluorescence intensity of both CDX2 and SOX2 compared with the DMSO control (Fig. 3A, B). This impairment was accompanied by a marked reduction in the total blastocyst cell number (Hoechst+) (Fig. 3C), resulting in significantly fewer TE (CDX2+) and ICM (SOX2+) cells (Fig. 3D, E).

**Fig. 3.**
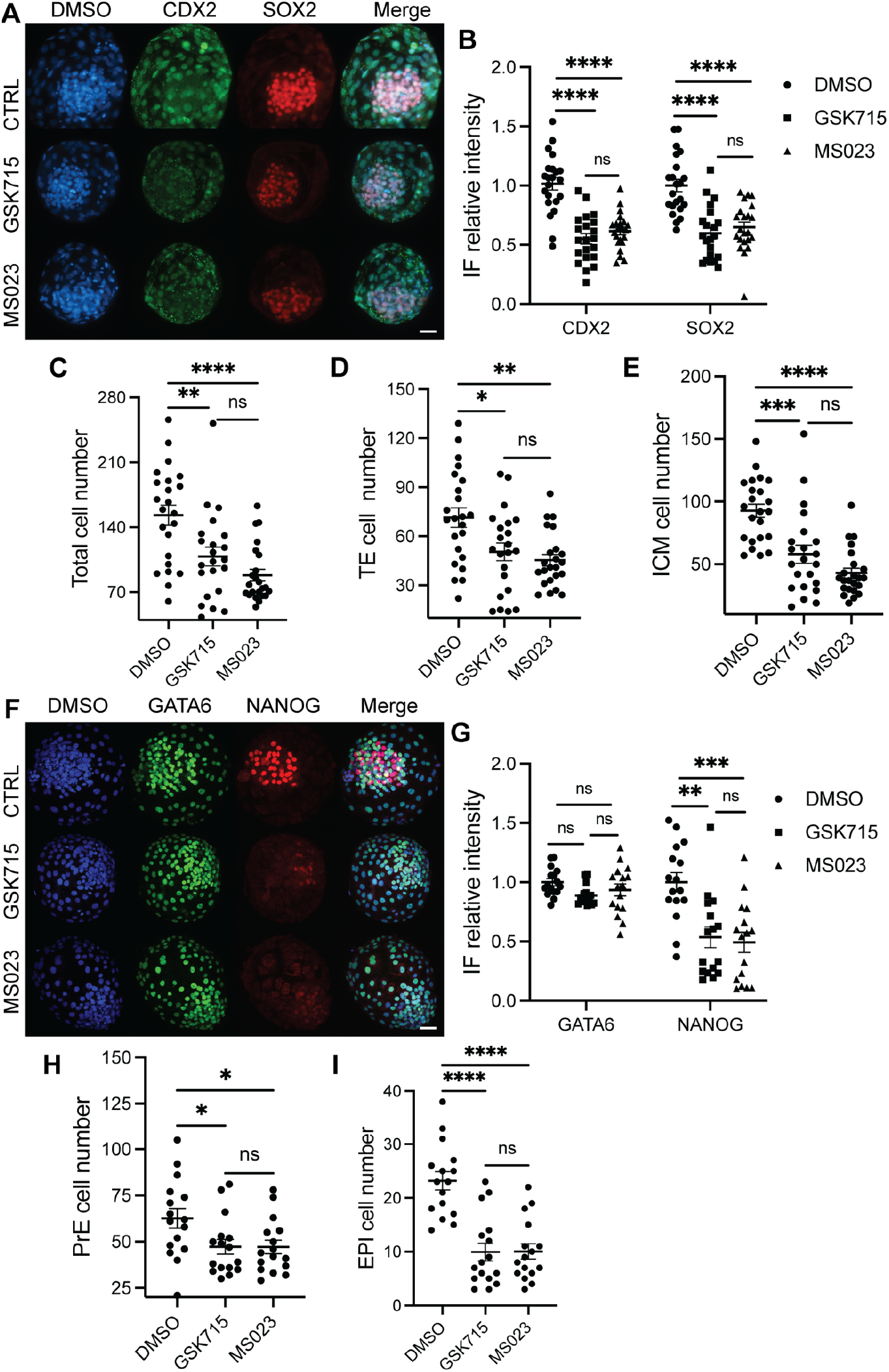
Inhibition of Type I PRMTs impairs lineage specification and cell allocation in cattle blastocysts. Immunofluorescence analysis of CDX2 (TE marker) and SOX2 (ICM marker) (A), and GATA6 (PrE marker) and NANOG (EPI marker) (F) in blastocysts treated with DMSO, GSK715, or MS023. Scale bar = 100 μm. Quantification of relative immunofluorescence (IF) intensity for CDX2 and SOX2 (B), and GATA6 and NANOG (G) in DMSO, GSK715 and MS023 groups. (C–E, H, I) Quantification of total cell number (C), TE cell number (D), ICM cell number (E), PrE cell number (H), and EPI cell number (I) per blastocyst in in DMSO, GSK715 and MS023 groups. n = 3 replicates, with 5–8 blastocysts per group per replicate. Each dot represents an individual blastocyst. n = 3 replicates, with 5-7 blastocysts per group per replicate. Data are presented as mean ± SEM. Asterisks in indicate significant differences (*P < 0.05, **P < 0.01, ***P < 0.001, ****P < 0.0001; ns, not significant).

We next evaluated the second cell fate decision by analysing the expression of the EPI marker NANOG and the PrE marker GATA6. Type I PRMT inhibition significantly reduced NANOG fluorescence intensity, whereas GATA6 expression remained unchanged (Fig. 3F, G). Nevertheless, the absolute numbers of both EPI (NANOG+) and PrE (GATA6+) cells were significantly decreased, reflecting impaired expansion of the inner cell mass (Fig. 3H, I).

Together, these findings demonstrate that Type I PRMT-mediated arginine methylation is required for both the first and second cell fate decisions during bovine preimplantation development, thereby supporting an essential role for ADMA-dependent epigenetic regulation in early lineage specification.

### PRMT inhibition impairs proliferation and TE-related transcription factor expression in cattle blastocysts

To determine whether the reduced blastocyst cell number resulted from impaired cell proliferation, we analysed the mitotic marker phospho-histone H3 (pH3S10). IF analysis revealed that the proportion of pH3S10-positive cells was significantly reduced following treatment with either GSK715 or MS023 compared with the DMSO control (Fig. 4A, B), indicating that inhibition of Type I PRMTs markedly suppresses mitotic activity during preimplantation development.

**Fig. 4.**
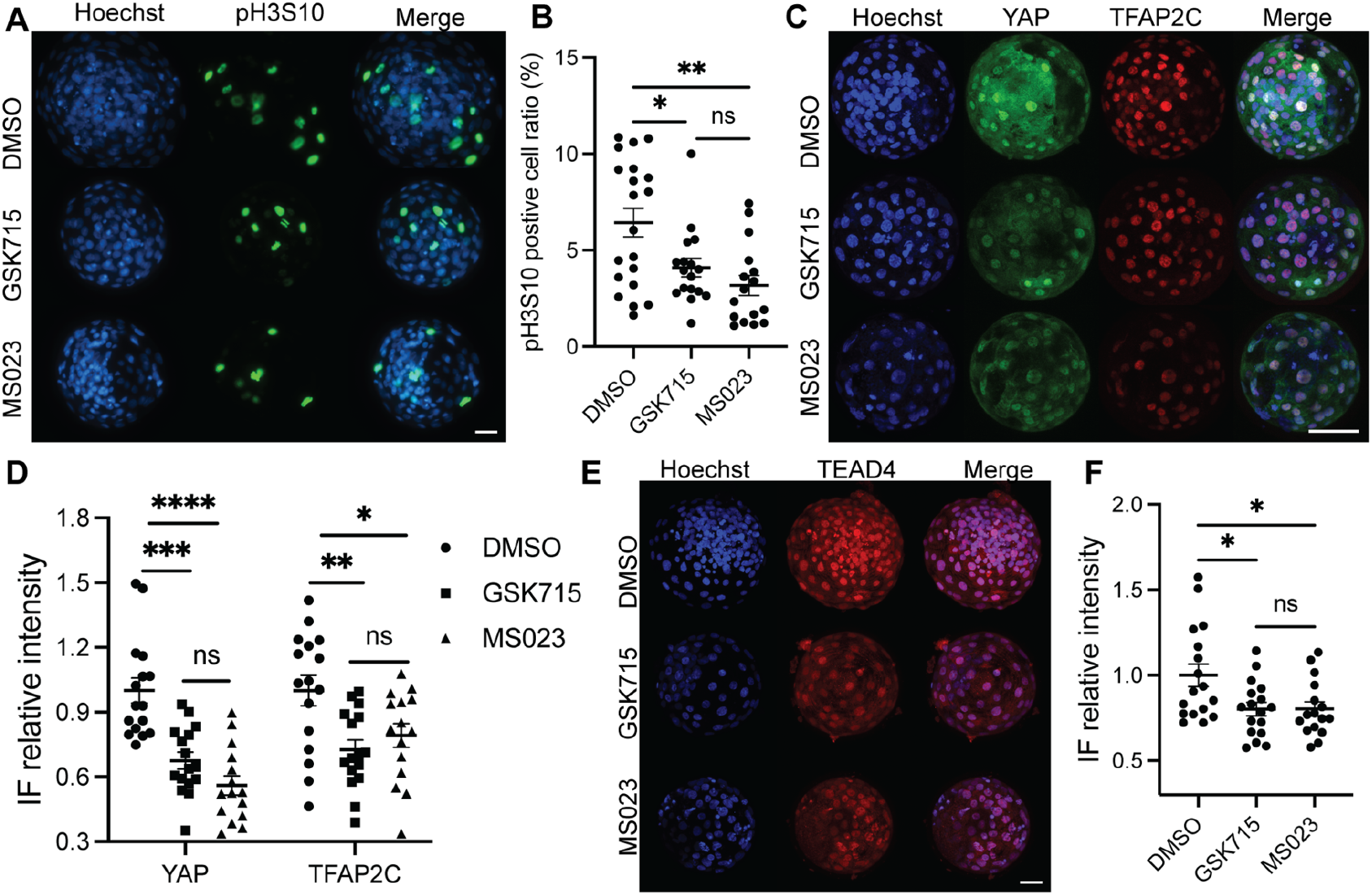
PRMT inhibition impairs proliferation and TE-related transcription factor expression in cattle blastocysts. Immunofluorescence analysis of pH3S10 (mitotic marker) (A), YAP and TFAP2C (C), and TEAD4 (E) in blastocysts treated with DMSO, GSK715, or MS023. Scale bars = 100 μm. Quantification of the percentage of pH3S10-positive cells (B), relative immunofluorescence (IF) intensity for YAP and TFAP2C (D), and relative IF intensity for TEAD4 (F) in DMSO, GSK715, and MS023 groups. n = 3 replicates, with 5–8 blastocysts per group per replicate. Each dot represents an individual blastocyst. n = 3 replicates, with 5-7 blastocysts per group per replicate. Data are presented as mean ± SEM. Asterisks in indicate significant differences (*P < 0.05, **P < 0.01, ****P < 0.0001; ns, not significant).

We next examined key components of the glucose-responsive Hippo signalling pathway [3]. IF analysis showed that the expression of YAP and TFAP2C was significantly reduced in embryos treated with either GSK715 or MS023 (Fig. 4C, D). Consistently, the expression of TEAD4, the DNA-binding partner of YAP required for TE specification, was also markedly decreased following Type I PRMT inhibition (Fig. 4E, F).

Together, these findings indicate that inhibition of Type I PRMT-mediated arginine methylation compromises TE specification through the combined suppression of embryonic cell proliferation and the Hippo-dependent YAP1–TEAD4–TFAP2C transcriptional programme.

### Inhibition of Type I PRMTs also impairs mouse TE differentiation

To determine whether Type I PRMT-mediated arginine methylation regulates lineage specification in a conserved manner in other mammalian embryos, we extended our analysis to mouse preimplantation embryos. Mouse embryos were treated with 1 μM MS023 and analysed at the blastocyst stage (Fig. 5A).

**Fig. 5.**
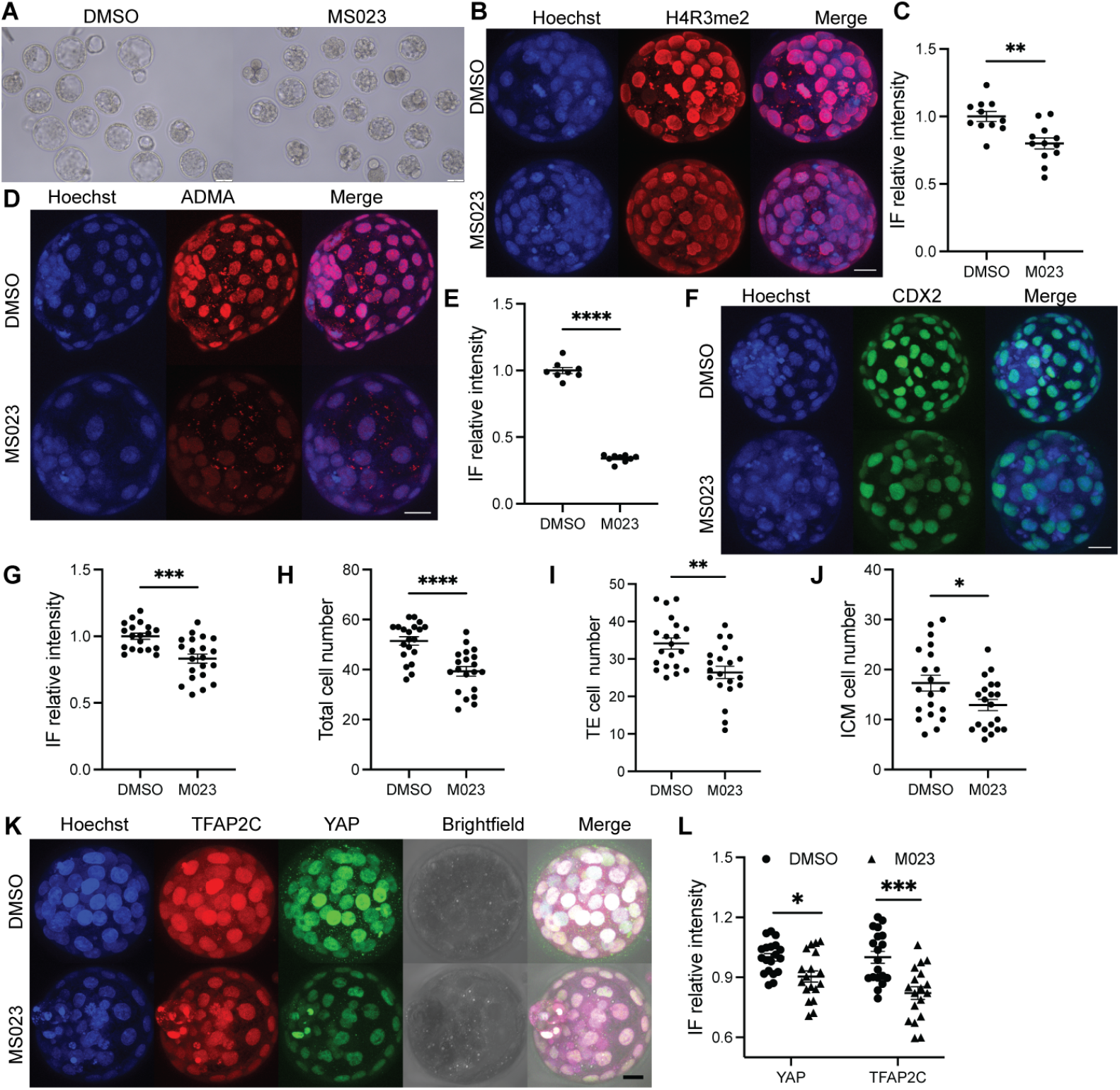
Inhibition of Type I PRMTs impairs mouse TE differentiation. (A) Representative bright-field images of mouse blastocysts treated with DMSO or MS023. Scale bar = 50 μm. (B, D, F, K) Immunofluorescence analysis of H4R3me2 (B), ADMA (D), CDX2 (TE marker) (F), and TFAP2C and YAP (K) in mouse blastocysts treated with DMSO or MS023. Scale bars = 100 μm. (C, E, G, L) Quantification of relative immunofluorescence (IF) intensity for H4R3me2 (C), ADMA (E), CDX2 (G), and YAP and TFAP2C (L) in DMSO and MS023 groups. (H–J) Quantification of total cell number (H), TE cell number (I), and ICM cell number (J) per mouse blastocyst in DMSO and MS023 groups. Data are presented as mean ± SEM. Each dot represents an individual blastocyst. n = 3 replicates, with 2– 4 blastocysts per group in each replicate. Asterisks indicate significant differences (*P < 0.05, **P < 0.01, ***P < 0.001, ****P < 0.0001).

Consistent with our findings in cattle, MS023 treatment markedly reduced H4R3me2a deposition (Fig. 5B, C) and global ADMA levels (Fig. 5D, E). We next examined the impact of Type I PRMT inhibition on the TE specification network. Expression of the key TE transcription factor CDX2 was significantly reduced following MS023 treatment (Fig. 5F, G). Likewise, the Hippo signalling components YAP and TFAP2C were also markedly downregulated (Fig. 5K, L). These molecular alterations were accompanied by impaired embryonic development. Type I PRMT inhibition significantly reduced the total cell number (DAPI+) of mouse blastocysts (Fig. 5H), resulting in marked decreases in both TE (CDX2+) (Fig. 5I) and ICM (CDX2-) (Fig. 5J) cell numbers. Together, these findings demonstrate that the Type I PRMT–ADMA–Hippo regulatory axis is conserved between cattle and mice.

## Discussion

In this study, we investigated the functional role of Type I PRMT-mediated ADMA in mammalian preimplantation cell fate determination. Our findings demonstrate that ADMA serves as a critical epigenetic prerequisite for proper early lineage specification and cell allocation. Furthermore, we established that the Type I PRMT–ADMA–Hippo regulatory axis is a highly conserved universal epigenetic mechanism driving these processes across mammalian species.

### Rationale for Pharmacological Inhibition over Genetic Ablation

To investigate the functional necessity of ADMA in early preimplantation development, we employed a pharmacological inhibition strategy rather than genetic ablation. The Type I PRMT family is composed of multiple distinct enzymes, all of which catalyze the formation of ADMA [6, 14]. Because these enzymes exhibit functional redundancy, knocking out or knocking down a single gene can trigger compensatory activity from other Type I members, preventing a complete blockage of ADMA synthesis [6, 14].

Furthermore, simultaneous multigene knockout of all Type I PRMTs in preimplantation mammalian embryos is technically prohibitive. To circumvent these limitations, we utilized GSK715 and MS023, two highly selective competitive inhibitors that explicitly target and block the active substrate-binding pockets of Type I PRMTs [12, 13]. Choosing two structurally distinct small molecules provides cross-validation, ensuring that the observed cellular and lineage defects result specifically from Type I PRMT inhibition rather than compound-specific off-target effects.

Among the Type I PRMT family, PRMT1 generates approximately 85% of total cellular ADMA [6]. Given its primary catalytic contribution, we specifically tracked PRMT1 protein distribution alongside its uniquely catalyzed histone modification, H4R3me2a (Fig. 1B-D). Pharmacological treatment with optimized doses of GSK715 or MS023 robustly reduced H4R3me2a and global ADMA (Fig. 2D, E), directly confirming that our inhibitor strategy effectively paralyzed PRMT1 activity on chromatin and disrupted global ADMA equilibrium.

Nevertheless, certain technical limitations of this study must be acknowledged. Due to the inherent scarcity and precious nature of preimplantation embryonic samples, we were unable to systematically profile all potential histone sites or non-histone substrates across every individual Type I PRMT. Consequently, it remains challenging to definitively determine whether the observed lineage specification failure is driven by the inhibition of a single specific Type I PRMT or by a systemic collapse in overall ADMA levels. Future work combining single-cell proteomic or target-specific mutational approaches will be essential to dissect the individual contributions of specific PRMT enzymes and their respective downstream substrates.

### Role of Type I PRMTs in ICM Specification and Pluripotency Maintenance

The profound impairment of ICM expansion and second lineage segregation (EPI/PrE) observed upon Type I PRMT inhibition aligns remarkably well with established roles of CARM1 in early mammalian embryogenesis. In mouse preimplantation embryos, CARM1-mediated asymmetric dimethylation of H3R26me2 serves as a primary epigenetic driver biasing blastomeres toward a pluripotent ICM fate [8, 9]. Blastomeres exhibiting elevated CARM1 activity display enhanced global transcription and direct up-regulation of core pluripotency factors, including Nanog, Sox2, and Sox21 [8, 9].

Conversely, genetic ablation or knockdown of robustly reduces H3R26me2 deposition, leading to a marked loss of NANOG-positive ICM cells and severe lineage misspecification [7]. Similarly, in porcine embryos, functional inactivation of CARM1 significantly suppresses H3R26me2 deposition and arrests blastocyst development due to a combined depletion of ICM and TE lineage allocation [15]. Given that both GSK715 and MS023 broadly target the shared catalytic pocket across Type I PRMTs [12, 13], the pronounced decline in ICM cell counts, NANOG expression, and EPI/PrE allocation (Fig. 3E-G) observed in cattle blastocysts is highly likely driven by the simultaneous enzymatic suppression of PRMT4 alongside other type I PRMTs.

### ADMA as a Potential Epigenetic Bridge Interlinking Glucose Metabolism, Hippo Signaling, and TE Specification

Beyond ICM specification, our findings uncover a vital role for Type I PRMT-mediated ADMA in TE differentiation. In mice, glucose availability acts as an essential metabolic driver of TE fate determination by triggering the YAP1–TEAD4–TFAP2C transcriptional axis to activate CDX2 [3]. Previous metabolomic profiling in bovine embryos revealed that fluctuating glucose levels directly alter intracellular SDMA and ADMA dynamics, suggesting an intricate crosstalk between glucose metabolism and arginine methylation [5]. Consistently, our data demonstrate that blocking ADMA synthesis via Type I PRMT inhibitors downregulates key core Hippo components (YAP, TEAD4, and TFAP2C) (Fig. 4C-F, 5K, 5L) and diminishes CDX2 expression (Fig. 4A, 4B, 5F, 5G). Mechanistically, this phenotypic overlap is echoed by single-embryo transcriptomic analyses in porcine embryos, where CARM1 knockdown directly downregulates key genes within the Hippo signaling pathway [15]. Furthermore, while CARM1 activity suppresses premature Cdx2 activation at early cleavage stages via Sox21, its activity is crucial at later stages to sustain proper cell division rates and TE cell numbers [7, 16]. Thus, our study strongly positions global ADMA deposition as a missing epigenetic bridge that converts upstream glucose metabolic cues into downstream Hippo pathway activation to drive TE specification.

### Mechanistic Limitations and Future Outlook

Notwithstanding these insights, a key limitation of the present study lies in dissecting the precise biochemical bridge connecting ADMA to Hippo pathway transducers.

Although our results demonstrate that ADMA depletion suppresses YAP, TEAD4, and TFAP2C levels, it remains to be determined whether ADMA regulates Hippo transducers directly via post-translational arginine methylation of YAP/TEAD4 proteins, or indirectly through chromatin accessibility and metabolic feedback loops. Future functional studies employing target-specific mutagenesis or quantitative methyl-proteomics will be required to define how ADMA mechanically fine-tunes Hippo pathway transducers to direct early lineage fate.

## Author contribution statement

R.S and P.B. conceived the project. J.Q. designed the experiments. J.Q. and Y.C. performed experiments and data analysis. J. Q. wrote the paper. R.S, P.B. and G.G. supervised the project.

## Acknowledgements

The authors thank Henry Leese for his helpful discussions.

---

**Supplementary Table S1.** Reagent information.

| Primary antibody | Concentration | Host | Supplier | CatLog NO. |
| --- | --- | --- | --- | --- |
| Anti-NANOG | 1:100 | Mouse | Thermo Fisher | 14-5768-82 |
| Anti-GATA6 | 1:1600 | Rabbit | Cell Signalling Technology | 5851T |
| Anti-SOX2 | 1:100 | Rat | Thermo Fisher | 14-9811-82 |
| Anti-ADMA | 1:200 | Rabbit | Cell Signalling Technology | 13522 |
| Anti-SDMA | 1:100 | Rabbit | Cell Signalling Technology | 13222 |
| Anti-MMA | 1:100 | Rabbit | Cell Signalling Technology | 8177S |
| Anti-CDX2 | 1:100 | Rabbit | Abcam | ab227201 |
| Anti-CDX2 | 1:100 | Mouse | Bio-Genex | AM392 |
| Anti-YAP | 1:100 | Rabbit | Cell Signalling Technology | 14074S |
| Anti-TFAP2C | 1:100 | Mouse | Santa Cruz | sc-12762 |
| Anti-TEAD4 | 1:100 | Mouse | Abcam | ab58310 |
| Anti-PRMT1 | 1:100 | Rabbit | Cell Signalling Technology | 2449S |
| Secondary antibody | Concentration | Host | Supplier | CatLog NO. |
| Alexa Fluor™ 488 | 1:100 | Goat anti-Mouse | Thermo Fisher | A-11001 |
| Alexa Fluor™ 594 | 1:100 | Goat anti-Rat | Thermo Fisher | A-11007 |
| Alexa Fluor™ 488 | 1:100 | Goat anti-Rabbit | Thermo Fisher | A-11008 |
| Alexa Fluor™ 594 | 1:100 | Goat anti-Rabbit | Thermo Fisher | A-11012 |
| Alexa Fluor™ 568 | 1:100 | Goat anti-Mouse | Thermo Fisher | A-11004 |
| Inhibitor | Concentration | Supplier | CatLog NO. |  |
| GSK3368715 | 10 µM | Cambridge bioscience | 1628925-77-8 |  |
| MS023 | 1 µM | Cambridge bioscience | 1831110-54-3 |  |

